# Optimization of skeletal protein preparation for LC-MS/MS sequencing yields additional coral skeletal proteins in *Stylophora pistillata*

**DOI:** 10.1101/2020.03.16.991273

**Authors:** Yanai Peled, Jeana Drake, Assaf Malik, Ricardo Almuly, Maya Lalzar, David Morgenstern, Tali Mass

## Abstract

Stony corals generate their calcium carbonate exoskeleton in a highly controlled biomineralization process mediated by a variety of macromolecules including proteins. Fully identifying and classifying these proteins is crucial to understanding their role in exoskeleton formation, yet no optimal method to purify and characterize the full suite of extracted coral skeletal proteins has been established and hence their complete composition remains obscure. Here, we tested four skeletal protein purification protocols using acetone precipitation and ultrafiltration dialysis filters to present a comprehensive scleractinian coral skeletal proteome. We identified a total of 60 proteins in the coral skeleton, 44 of which were not present in previously published stony coral skeletal proteomes. Extracted protein purification protocols carried out in this study revealed that no one method captures all proteins and each protocol revealed a unique set of method-exclusive proteins. To better understand the general mechanism of skeletal protein transportation, we further examined the proteins’ gene ontology, transmembrane domains, and signal peptides. We found that transmembrane domain proteins and signal peptide secretion pathways, by themselves, could not explain the transportation of proteins to the skeleton. We therefore propose that some proteins are transported to the skeleton via non-traditional secretion pathways.

## Introduction

Scleractinian corals, or stony corals, are the most prolific biomineralizers in phylum Cnidaria [1]. They are a key component of shallow-water tropical reefs, often forming massive structures that serve as the foundation of an ecosystem which hosts some of the more biodiverse communities on the planet [2, 3]. They are amongst the oldest biomineralizing metazoans, producing calcium carbonate (CaCO_3_) exoskeletons in the form of aragonite through biologically-directed mechanisms [4, 5], which makes up ≥95% of the entire skeletal mass, the remainder of which is the skeletal organic matrix (SOM) [6-8].

Biomineralization refers to the ability of a living organism to selectively exploit elements from its surrounding environment to build a biologically functioning crystalline structure [9, 10]. Biomineralizers can be found throughout all kingdoms of life, from bacteria [e.x. 11], to algae [e.x., 12], mollusks [e.x., 13], corals [e.x., 10] and mammals [e.x., 14]. The minerals formed through this process differ in structure from their non-biological counterparts and their formation is mediated by a variety of organic molecules (the SOM), which have been intensively studied since the 1960’s in diverse organisms [reviewed by 10]. The SOM consists of proteins, lipids, and polysaccharides, which are not present in the abiotic mineral form [reviewed by 15]. SOM proteins (SOMPs) embedded within the mineral are hypothesized to serve as a framework for crystal nucleation [reviewed by 16]. The proteins involved in the process of skeletal biomineralization have been described and studied extensively in echinoderms [e.x., 17, 18] [and reviewed by 19, 20], mollusks [e.x., 21, 22-24] and mammals [e.x., 25, 26, 27], among others [28]. At present, the best described SOMP complex is of mammalian bone and teeth [15].

In stony corals, the most current knowledge of SOMPs is limited and based on intraskeletal protein extraction [29-33]. It has been suggested that coral SOMPs aid in the molecular processes of crystallization as well as in the development and strengthening of the minerals ([34-37] among others). The constant advancement of mass spectrometry technology has broadened our capability to identify many proteins in skeletal extracts, even those proteins in low abundance [38]. However, this technology is sensitive to contamination by organic matter remnants from soft tissue and cell debris from the study organism, and the little-addressed issue remains of contamination by researchers during the protein extraction, preparation, and sequencing steps [39-41]. Upon extraction, the SOMPs are usually divided into two fractions: soluble and insoluble matrix proteins (SSOM and ISOM respectively), based on their solubility in the acid of choice or in water [22, 31-33, 42]. While some past attention has been directed towards the soluble fraction [30, 43, 44], Pereira-Mouriès et al. [45] showed that, in the bivalve *Pinctada maxima*, the classification of SSOM and ISOM is misleading and that both fractions share common features.

Furthermore, Goffredo et al. [42] found in the stony coral *Balanophyllia europaea* that both fractions consist of the same macromolecules; they associated the degree of solubility to the difference in cross-linking. They also showed that each solubility fraction has a different influence on calcium carbonate crystal morphology, aggregation, and polymorphism in-vitro. In contrast, Ramos-Silva et al. [32] observed a different SOMP composition between solubility fractions in the scleractinian coral *Acropora millepora*,. Out of 36 SOMPs, only two were found exclusively in the soluble fraction and twelve were exclusive to the insoluble fraction. These examples demonstrate the attempts to attribute different properties to the two fractions, but the data remain inconclusive.

To date, three major coral skeletal proteomes have been published [31-33] with each proteome consisting of 30-40 proteins. Of the 30 proteins sequenced from *A. digitifera* skeleton, 26 were also detected in *A. millepora* skeleton [32, 33]. They consist mostly of either transmembrane (TM) domain proteins or secretory proteins [33]. However, only 12 of the proteins identified in *A. millepora* skeleton matched those found in *S. pistillata* skeleton [31, 32]. In *A. millepora*, 11 TM domain-containing proteins were identified, as well as two proteases that were not detected in *S. pistillata* [32]. The authors suggested that the proteases’ role is in cleaving the extracellular domain of TM proteins and incorporating them into the skeleton.

The coral skeletal proteomes published to date reveal an overlap of several detected proteins, but at least 10 proteins from each species appear to be unique. It is currently unknown if this is truly due to species-specific gene expression and protein localization or to methods in extracting, purifying, and sequencing the proteins. In this study we analyzed several methods for extracted protein purification to increase the detection of the full suite of SOMPs from cleaned coral skeleton powder. We show that the use of acetone precipitation versus centrifugal filter washing, and the degree to which each purification method is performed, affects the numbers and types of proteins that can be sequenced by mass spectrometry. Further, we suggest that there is no one ‘best’ method for coral skeletal protein purification to capture all SOMPs such that future research projects may need to utilize several preparation methods to detect the full breadth of proteins embedded in coral skeleton.

## Methods

### Sample collection and preparation for protein extraction

The hermatypic coral *Stylophora pistillata* (Esper, 1797) was collected under a special permit from the Israeli Natural Parks Authority in the waters in front of the H. Steinitz Marine Biology Laboratory, Eilat, Israel, Red Sea (29°30 N, 34°56 E), using SCUBA diving.

We fragmented one *S. pistillata* colony into small pieces, approximately 2×2 cm, with a diamond band saw. Coral fragments were transferred to 50-ml Falcon brand conical vials (Falcon tubes) and oxidized with 20 mL 1:1 of 30% H_2_O_2_: 3% NaClO solution for 1 hour, during which 1.5 mL of 3% NaClO solution were gently added to the tubes every 20 minutes and continued the incubation overnight at room temperature following modified methods of Stoll et al. [46]. Fragments were washed five times with ultra-pure water for one minute each time and dried at 60 C° overnight. We crushed the cleaned fragments to ≤63 µm diameter with a mortar and pestle. Skeleton powder, in sterile Falcon tubes, was then oxidized and washed in ultra-pure water three more times (i.e., four complete rounds of oxidative cleaning) to ensure that no organic residue remained on the skeletal grains. In each cycle, the removal of the oxidizing or wash solution was performed by centrifugation at 5,000 x g for 3 min at 4 C°. Cleaned skeletal powder was then dried overnight at 60 C°. We carried out all the described processes in a laminar flow biological hood (apart from oven drying) with all preparation tools and surfaces bleached to avoid contamination.

To monitor the removal of proteins from the skeletal powder, we checked the cleaning efficiency under SEM after the fourth oxidative cleaning. Samples were sputter-coated with 4 nm gold prior to examination using a ZEISS Sigma TM scanning electron microscope with in-lens detector (5kV, WD = 5-7mm) (SI Figure 1a,b) [17, 47]. In addition, we sonicated the cleaned powder at 4°C in filter-sterilized phosphate buffered saline (PBS, pH 7.4) for 30 minutes, pelleted the powder at 5,000 x *g* for 3 minutes at 4°C, concentrated the supernatant on a 3-kDa cutoff centrifugal filter unit (Amicon) and loaded samples of supernatant on a 8-16% SDS-PAGE TGX Stain-Free gels (Bio-Rad) (SI Figure 1c).

### Extraction and purification of skeletal proteins

We used four samples of approximately 1.3 g cleaned skeleton powder each to test four protein purification protocols as described in Figure 1. All samples were decalcified in 0.5 M acetic acid (30 ml acid/g cleaned skeleton powder) in Falcon tubes while rotating the tubes at room temperature for 3 hours. Samples were then centrifuged at 5,000 x *g* for 5 min at 4°C and supernatant was transferred to a new tube and stored at 4°C. We continued the decalcification of the undissolved pellets with a second volume of 0.5 M acetic acid and allowed decalcification to proceed to completion, at which point the pH of the solution was measured at 5.5-6.5. We then combined both decalcification rounds (70 ml total) for each sample containing both the acid-soluble and -insoluble fractions, froze the total volumes at -80°C, and dried them by overnight lyophilization. The dried pellets were stored at *-*80°C until further processing. Because washes of the extracted proteins can reduce the representation of proteins, in this study we examined two protein concentrating and cleaning methods similar to those performed for previously published coral skeletal proteomes; (i) centrifugation ultrafiltration (CF methods [32]) and (ii) acetone precipitation (ACT methods, [31]) (Figure 1; SI Table 1).

**Figure 1.**
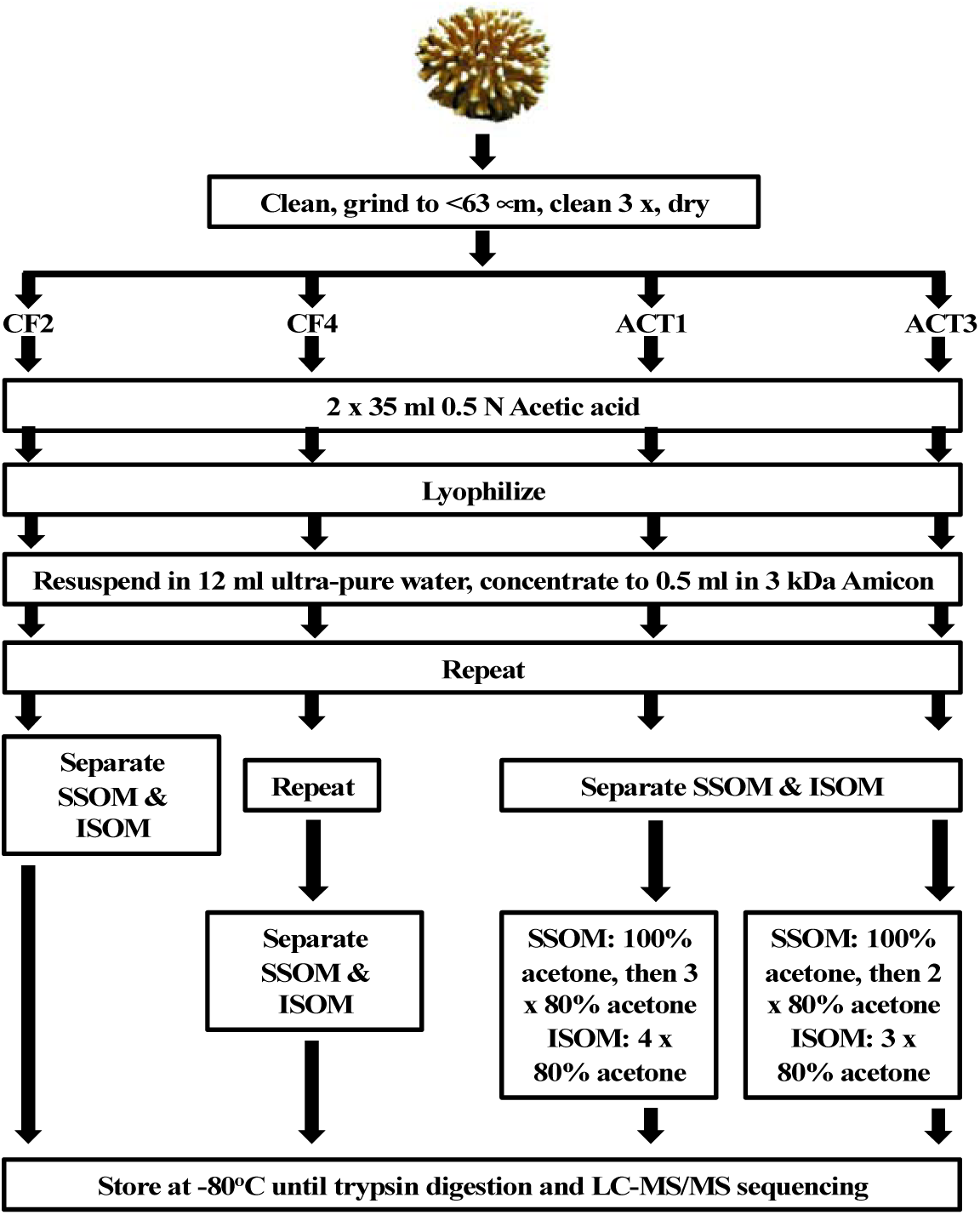
Workflow of methods for skeletal protein purification and concentration after extraction in acetic acid. CF – Centrifugation filtration methods; ACT – acetone precipitation methods; SSOM –soluble skeletal organic matrix; ISOM –insoluble skeletal organic matrix.

To desalt and concentrate the extracted SOM, the lyophilized pellets of all samples were re-suspended in 12 ml MilliQ water and the proteins were concentrated on 3 kDa cutoff Amicon® Ultra 15 centrifugal filter units (Merck-Millipore) 5,000 x *g* at 4°C to reach a final volume of 0.5 ml. This process was repeated once. At this stage, we observed a water-insoluble pellet (ISOM) in all samples, which for samples CF2, ACT1, and ACT3 was pelleted from the water-soluble fraction (SSOM) by centrifugation at 5,000 x g for 5 min at 4°C. This initial wash, concentration, and solubility fractionation protocol was the terminal step for centrifugal filter sample CF2. Sample CF4 was resuspended one more time in MilliQ and concentrated before separation of SSOM and ISOM as above. For acetone precipitation samples ACT1 and ACT3, the two rounds of desalting and concentration were followed upon with successive washes of the SSOM and ISOM in acetone. To sample ACT1 SSOM was added 2 ml 100% ice cold acetone. The sample was vortexed for 10 seconds, incubated at -20°C for 30 minutes, and centrifuged at 4,300 x g for 30 min at 4°C. The resulting pellet was washed three more times with 2 mL of ice cold 80% acetone. The ISOM fraction was washed four times with 80% acetone. Both solubility fractions of sample ACT3 were treated as in ACT1 but with one less washing step of each fraction. All fractions across all purification treatments were stored at -80°C. An aliquot of each fraction was analyzed by SDS-PAGE on a 8-16% SDS-PAGE TGX Stain-Free gels with subsequent silver staining and displays the commonly-observed smearing of extracted biomineral proteins with minimal observable banding (SI Figure 2).

### LC MS\MS

*S. pistillata* skeletal protein samples were dissolved in 5% SDS and digested with trypsin using the S-trap method overnight at room temperature. We analyzed the resulting peptides using a nanoflow ultra-performance liquid chromatograph (nanoAcquity) coupled to a high resolution, high mass accuracy mass spectrometer (Fusion Lumos). The sample was trapped on a Symmetry C18 0.18*20mm trap column (Waters, Inc) and separated on a HSS T3 0.075*250 mm column (Waters, Inc.) using a gradient of 4-28% (80% acetonitrile, 0.1% Formic acid) for 150 minutes. Spray voltage was set to +2kV. The data were acquired in the Fusion Lumos using a Top Speed Data-Dependent Acquisition method using a cycle time of 3 s. An MS1 scan was performed in the Orbitrap at 120,000 resolution with a maximum injection time of 60 ms. The data were scanned between 300-1800 m/z. MS2 was selected using a monoisotopic precursor selection set to peptides, peptide charge states set to +2-+8 and dynamic exclusion set to 30 s. MS2 was performed using HCD fragmentation scanned in the Orbitrap, with the first mass set to 130 m/z at a resolution of 15,000. Maximum injection time was set to 60 ms with automatic gain control of 5×10^−4^ ions as a fill target. The resulting data were searched against the NCBI *Stylophora pistillata* protein database using the Byonic search engine (Protein Metrics Inc.) – the first search was carried out without any false discovery rate (FDR) filtering, to generate a focused database for a second search. The second search was set to 1% FDR, allowing fixed carbamidomethylation on C and variable oxidation on MW, deamidation on NQ and protein N-terminal acetylation. The mass spectrometry proteomics data have been deposited to the ProteomeXchange Consortium via the Pride partner repository [48], under the dataset identifier PXD017891; reviewer login username and password have been provided to the reviewers. Data will be made publicly available without a required login in ProteomeXchange upon publication of the manuscript.

### Data sorting

We used the *S. pistillata* genome database as a reference peptide database for the mass spectrometry analysis [49] (NCBI BioProjects PRJNA281535 and PRJNA415215) appended with known NCBI *S. pistillata* skeletal proteins[e.x., 31, 50] *S. pistillata* carbonic anhydrase, ACE95141.1). We also included a common contaminants database. Despite this inclusion of a contaminants database, several proteins likely of human origin were sequenced and attributed to *S. pistillata*. To filter out these potential contaminants from our final list of coral-specific proteins, we BLASTed all sequences against the ‘Primates’ database in NCBI using Blast2GO. We then examined NCBI-generated sequence alignments of coral versus *Homo sapiens* proteins with e-values lower than e^−50^ and percent mean similarity greater than 50%, all sequences with e-values lower than e^−100^, and all sequences with percent similarity greater than 80%, and removed from our final list of coral proteins any sequences with three or more peptides each of seven or more amino acids in length that were identical between *S. pistillata* and humans.

All remaining coral-specific proteins identified by the LC MS\MS analysis were filtered to those with at least two significant peptides or at least one significant peptide with at least 10 spectra and an identification score of 250 or greater. Skeletal proteins were first sorted by fractions and methods (i.e., SSOM and ISOM for each purification method). Next, we sorted all skeletal proteins by their gene ontology (GO) terms [51, 52]. Finally, we grouped all by terms of interest: protein modification, transmembrane, and ECM; membrane processing; metal binding and vesicular.

Proteins detected by LC-MS/MS were annotated using the Trinotate pipeline which relies on both Pfam and UniProt data [53]. GO terms and Pfam annotations were assigned to *Stylophora pistillata* predicted proteins and transcripts using Trinotate 3.0.1 (https://github.com/Trinotate). Transmembrane regions were predicted using the TMHMM server v2.0 [54]. Signal peptides on the N-termini of proteins were predicted using SignalP 5.0 [55, 56]. Attachment of proteins to the exterior of the cell membrane by glycosylphosphatidylinositol (GPI) anchors was predicted using PredGPI [57]. These computational analyses used program default settings and cutoffs. Completeness of protein sequences was determined by comparing all returned coral proteins to the *Acropora digitifera* genome [58]; NCBI RefSeq assembly GCF_000222465.1). Detection of various proteins across solubility and protocol fractions was visualized in Venny 2.1 online software.

To examine the conserved coral biomineralization proteins, we determined orthologous biomineralization genes, known from skeletal proteomic analysis, across coral taxa. First, we estimated orthology relationships between all non-redundant genes of selected metazoa species using OrthoFinder. We included all genes of all Cnidaria species with known genome-based annotations. OrthoFinder generates orthology groups (Orthogroups) based on normalized reciprocal best BLAST hits’ bit scores [59], and then estimates orthologues genes pairs within Orthogroups [60]. We then selected all pairs of *Acropora* spp. orthologs to *S. pistillata* (1:1, 1:many, many:many relationships) ([31-33], this study). From these pairs we further selected *S. pistillata* spectra-based identified proteins, or skeletal proteins known in the literature. Since not all skeletal protein annotations from the literature were included in our reference OrthoFinder proteome datasets, we further found their best matches in the reference OrthoFinder proteome using BLASTP.

## Results

After extensive cleaning of the powdered skeleton and acid-extraction of embedded organic matter we identified in total 60 coral-specific proteins meeting our criteria in *S. pistillata* skeleton as predicted by the species genome [49] (Table 1, SI Table 2). Trinotate [53, 61] returned GO annotations [51] for all of these proteins, although many remain uncharacterized (SI Table 3).

**Table 1:** 60 coral skeletal proteins detected by LC-MS/MS across all treatments and solubility fractions. Proteins are listed in order of accession number. Geno ontology categorization is represented as ^a^ECM/transmembrane and protein modification, ^b^membrane processing, and ^c^vesicle/secretion and metal binding.

| Stylophora pistillata genome hit |  |  | Solubility Fraction |  |  |  |
| --- | --- | --- | --- | --- | --- | --- |
| gi Number | Accession Number | Annotation | Best score | Go Categorization | ASM | AIM |
| 1263115389 | PFX12726.1 | Retrovirus-related Pol polyprotein from transposon 17.6 [Stylophora pistillata] | 340.80 | - |  | x |
| 1263115517 | PFX12813.1 | hypothetical protein AWC38_SpisGene23165 [Stylophora pistillata] | 337.40 | - |  | x |
| 1263116757 | PFX13778.1 | Sacsin [Stylophora pistillata] | 267.80 | - |  | x |
| 1263117261 | PFX14205.1 | Proto-oncogene tyrosine-protein kinase receptor Ret, partial [Stylophora pistillata] | 336.70 | a,c |  | x |
| 1263119001 | PFX15740.1 | Protein FAM208A [Stylophora pistillata] | 373.20 | - | x | x |
| 1263119725 | PFX16398.1 | hypothetical protein AWC38_SpisGene19330 [Stylophora pistillata] | 273.10 | - |  | x |
| 1263122270 | PFX18785.1 | Mucin-4 [Stylophora pistillata] | 449.50 | a,c |  | x |
| 1263130352 | PFX26597.1 | Complement C3 [Stylophora pistillata] | 283.10 | a,b,c |  | x |
| 1263130510 | PFX26751.1 | Transmembrane protease serine 9 [Stylophora pistillata] | 244.20 | a |  | x |
| 1263131615 | PFX27832.1 | Poly [ADP-ribose] polymerase 11 [Stylophora pistillata] | 320.10 | c |  | x |
| 1263134664 | PFX30831.1 | hypothetical protein AWC38_SpisGene4366 [Stylophora pistillata] | 322.20 | - |  | x |
| 1263134737 | PFX30903.1 | hypothetical protein AWC38_SpisGene4292 [Stylophora pistillata] | 548.90 | - |  | x |
| 1263135656 | PFX31810.1 | Nidogen-2 [Stylophora pistillata] | 286.20 | a,c |  | x |
| 1270028962 | XP_022783323.1 | phosphopantothenoyl cysteine decarboxylase subunit VHS3-like [Stylophora pistillata] | 406.60 | c |  | x |
| 1270029544 | XP_022786582.1 | synapsin-2-like isoform X2 [Stylophora pistillata] | 320.50 | c | x | x |
| 1270031975 | XP_022799089.1 | CUB domain-containing protein-like isoform X2 [Stylophora pistillata] | 699.70 | - |  | x |
| 1270032917 | XP_022804012.1 | EGF and laminin G domain-containing protein-like [Stylophora pistillata] | 384.10 | a |  | x |
| 1270036141 | XP_022779720.1 | vitellogenin-like [Stylophora pistillata] | 416.00 | b,c |  | x |
| 1270037242 | XP_022780303.1 | LOW QUALITY PROTEIN: uncharacterized protein LOC111321626 [Stylophora pistillata] | 353.60 | a,b,c | x |  |
| 1270037961 | XP_022780690.1 | skeletal aspartic acid-rich protein 2-like [Stylophora pistillata] | 723.60 | a |  | x |
| 1270037969 | XP_022780694.1 | CUB and peptidase domain-containing protein 2-like [Stylophora pistillata] | 285.80 | a | x | x |
| 1270041141 | XP_022782398.1 | skeletal aspartic acid-rich protein 1-like [Stylophora pistillata] | 709.80 | - | x | x |
| 1270042352 | XP_022783044.1 | uncharacterized protein LOC111323869 [Stylophora pistillata] | 283.60 | - |  | x |
| 1270043038 | XP_022783415.1 | coadhesin-like isoform X3 [Stylophora pistillata] | 411.20 | a |  | x |
| 1270044042 | XP_022783952.1 | collagenase 3-like [Stylophora pistillata] | 262.90 | a,c |  | x |
| 1270045301 | XP_022784623.1 | cation channel sperm-associated protein subunit beta-like [Stylophora pistillata] | 328.40 | a,c |  | x |
| 1270049598 | XP_022786918.1 | major yolk protein-like isoform X2 [Stylophora pistillata] | 476.50 | a,c |  | x |
| 1270052030 | XP_022788227.1 | hephaestin-like protein [Stylophora pistillata] | 539.80 | a,c |  | x |
| 1270052971 | XP_022788730.1 | chymotrypsin-like elastase family member 1 [Stylophora pistillata] | 264.80 | a,c |  | x |
| 1270054595 | XP_022789591.1 | endothelin-converting enzyme 1-like isoform X2 [Stylophora pistillata] | 94.10 | a,b,c |  | x |
| 1270055217 | XP_022789932.1 | MAGUK p55 subfamily member 7-like [Stylophora pistillata] | 313.70 | a |  | x |
| 1270056169 | XP_022790441.1 | PHD finger protein 21A-like [Stylophora pistillata] | 277.90 | c |  | x |
| 1270059479 | XP_022792212.1 | ras-like protein 3 [Stylophora pistillata] | 289.90 | a,c |  | x |
| 1270063049 | XP_022794122.1 | galaxin-like isoform X2 [Stylophora pistillata] | 309.40 | - |  | x |
| 1270063475 | XP_022794351.1 | mammalian ependymin-related protein 1-like [Stylophora pistillata] | 324.80 | a,c |  | x |
| 1270064196 | XP_022794736.1 | MAM and LDL-receptor class A domain-containing protein 2-like [Stylophora pistillata] | 472.20 | - |  | x |
| 1270068394 | XP_022796981.1 | uncharacterized skeletal organic matrix protein 8-like [Stylophora pistillata] | 664.40 | - |  | x |
| 1270068396 | XP_022796982.1 | uncharacterized protein LOC111335364 [Stylophora pistillata] | 609.30 | - |  | x |
| 1270071953 | XP_022798902.1 | low-density lipoprotein receptor-related protein 8-like [Stylophora pistillata] | 257.10 | c |  | x |
| 1270073139 | XP_022799541.1 | uncharacterized protein LOC111337489 [Stylophora pistillata] | 461.10 | b |  | x |
| 1270080553 | XP_022803524.1 | digestive cysteine proteinase 1-like [Stylophora pistillata] | 262.20 | a,c |  | x |
| 1270081088 | XP_022803808.1 | deleted in malignant brain tumors 1 protein-like [Stylophora pistillata] | 344.70 | c |  | x |
| 1270081207 | XP_022803872.1 | spore wall protein 2-like isoform X3 [Stylophora pistillata] | 338.60 | - |  | x |
| 1270081241 | XP_022803894.1 | uncharacterized protein LOC111341206 [Stylophora pistillata] | 390.70 | b |  | x |
| 1270082891 | XP_022804785.1 | thioredoxin reductase 1, cytoplasmic-like [Stylophora pistillata] | 337.00 | c |  | x |
| 1270084203 | XP_022805470.1 | uncharacterized protein LOC111342641 [Stylophora pistillata] | 256.10 | - |  | x |
| 1270085816 | XP_022806326.1 | ZP domain-containing protein-like [Stylophora pistillata] | 393.90 | a |  | x |
| 1270086467 | XP_022806664.1 | protein lingerer-like [Stylophora pistillata] | 250.10 | b,c |  | x |
| 1270086954 | XP_022806928.1 | SLIT-ROBO Rho GTPase-activating protein 1-like [Stylophora pistillata] | 309.10 | - |  | x |
| 1270087345 | XP_022807143.1 | condensin-2 complex subunit D3-like [Stylophora pistillata] | 278.10 | c |  | x |
| 1270087556 | XP_022807256.1 | uncharacterized protein LOC111344300 [Stylophora pistillata] | 280.40 | a,c |  | x |
| 1270088573 | XP_022807807.1 | uncharacterized protein LOC111344812 [Stylophora pistillata] | 365.30 | a,b,c |  | x |
| 1270089244 | XP_022808163.1 | uncharacterized protein LOC111345150 [Stylophora pistillata] | 310.20 | - |  | x |
| 1270090022 | XP_022808576.1 | uncharacterized protein LOC111345553 isoform X2 [Stylophora pistillata] | 320.10 | a,c |  | x |
| 1270091315 | XP_022809269.1 | microtubule-associated tumor suppressor 1 homolog isoform X1 [Stylophora pistillata] | 259.20 | a,c |  | x |
| 1270091317 | XP_022809270.1 | microtubule-associated tumor suppressor 1 homolog isoform X2 [Stylophora pistillata] | 256.40 | - |  | x |
| 1270093788 | XP_022810585.1 | von Willebrand factor D and EGF domain-containing protein-like, partial [Stylophora pistillata] | 298.80 | - |  | x |
| 1270095516 | XP_022778254.1 | uncharacterized protein LOC111319781 [Stylophora pistillata] | 250.70 | c |  | x |
| 1270095572 | XP_022778283.1 | uncharacterized protein LOC111319816, partial [Stylophora pistillata] | 508.10 | - | x | x |
| 190710633 | ACE95141.1 | carbonic anhydrase [Stylophora pistillata] | 495.20 | a,c | x | x |

In order to evaluate the efficacy and improve current methods for stony coral skeletal protein purification, we examined four different protocols; two centrifugation ultrafiltration filters (CF) and two further acetone precipitation (ACT) protocols. Proteomes of the four methodologies differed in composition and variety (Figure 2). Combining results of all CF fractions identified 52 coral-specific proteins while combined ACT protocols yielded 13 such proteins (Figure 2A). Moreover, redundancy between methodologies was low. Only 8.3% of the proteins overlapped between methods while 78.3% and 13.3% of the proteins were exclusive to combined CF and combined ACT fraction data, respectively.

**Figure 2:**
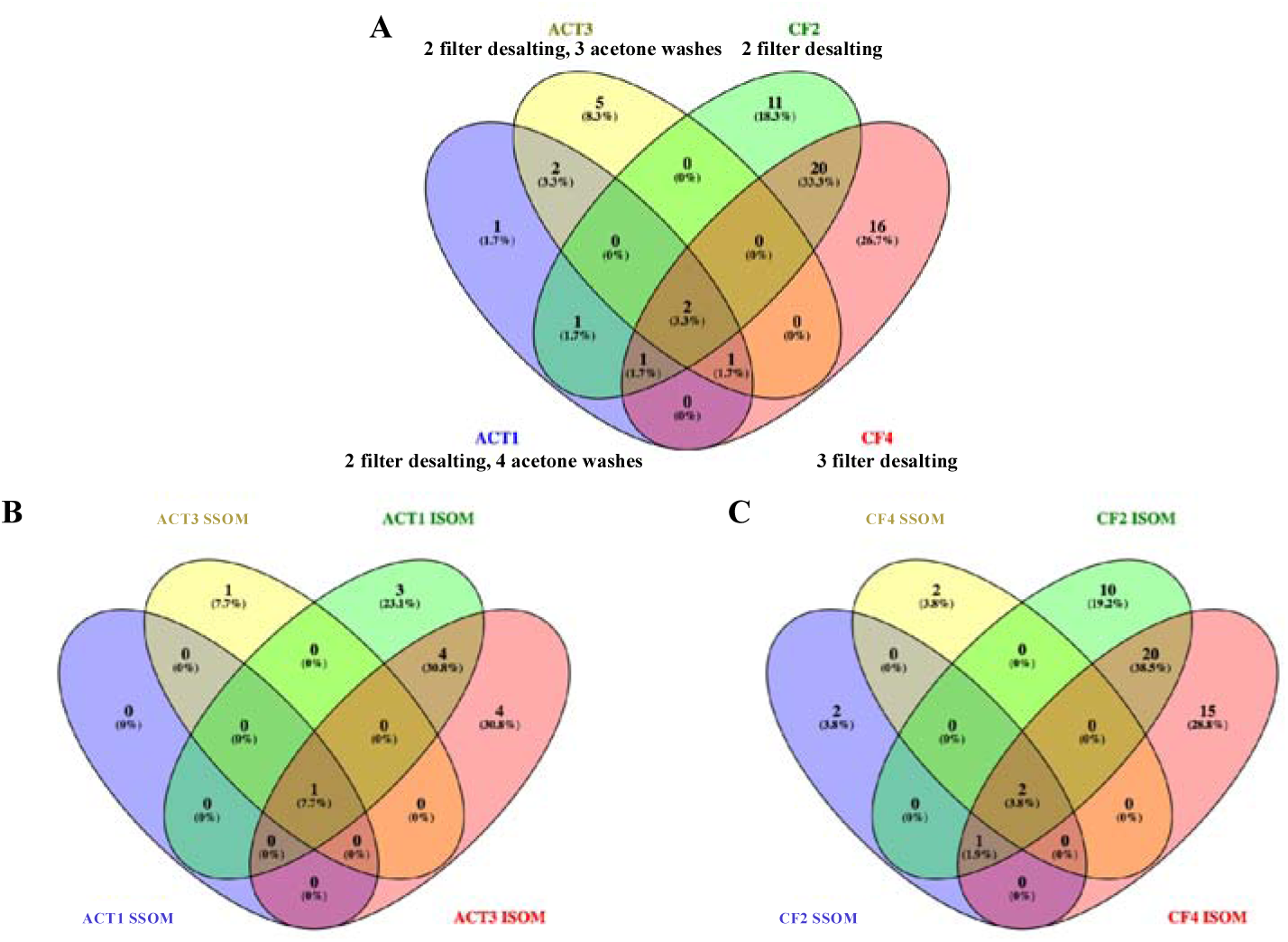
Distribution of proteins numbers according of the purification methods. Distribution of all proteins (SOM and ISOM combined) by purification methods (A). SOM and ISOM distribution by acetone precipitation methods (ACT; B). SOM and ISOM distribution by centrifugation and ultrafiltration method (CF; C). Note that CF methods yielded more proteins in total. ISOM Insoluble Skeletal Organic Matrix; SOM Skeletal Organic Matrix.

To evaluate the purification efficiency of each protocol, we first compared the number of proteins detected in each method (e,x,, ACT1 vs ACT 3 and CF2 vs CF4). Of the 8 proteins found only in samples from the acetone wash protocols (ACT 1 and ACT 3), one was observed only in ACT1 while five were observed only ACT3 (Figure 2, SI Table 2); both ACT1 solubility fractions had one more wash step than did those in ACT3. In contrast, of the 47 proteins observed only in CF samples, 12 were found only in CF2, which went through two filter centrifugation steps, while 16 were found only in CF4, which went through a third filter centrifugation step; the remaining 19 were found using both CF protocols. Further, only two proteins were observed in all four purification fractions in at least one solubility form: CARP4/SAARP1 and synapsin 2-like. Two additional proteins were observed in three of the four protocols: STPCA2 and an uncharacterized protein.

We next examined the difference in protein composition obtained from the solubility fractions SSOM versus ISOM. We identified different distributions of proteins in the SSOM vs. ISOM in both purification methods (Figures 2B & C). Notably, when combining all methods, more proteins were identified in the ISOM compared to the SSOM. A total of 57 coral proteins were identified in the ISOM fraction compared to a total of 8 in the SSOM fraction (SI Table 2), of which 5 (8.3%) were identify both in SSOM and ISOM.

### Our data compared to other coral skeletal proteomes

Since our analysis yielded a large amount of new skeletal proteins, we also compared our results with the three previously published proteomes of *S. pistillata, A. digitifera* and *A. millepora* [31-33]. Out of our entire identified skeletal proteome containing 60 proteins, using OrthoFinder and BLASTP, only 16 were found to be similar to proteins identified in these studies. Yet, this proportion of overlap (16 out of 60) is significantly greater than the expected proportion by chance, since the proportion of known skeletal matrix proteins in the reference coral proteomes is extremely small (less than ∼0.2%). Seven proteins were found to overlap all four proteomes: a coadhesin-like protein, an EGF and laminin G domain-containing protein, a hypothetical protein, a MAM and LDL-receptor class A domain-containing protein, a mucin, aspartic acid-rich protein 2-like, and a ZP domain-containing protein (Table 2).

**Table 2.**
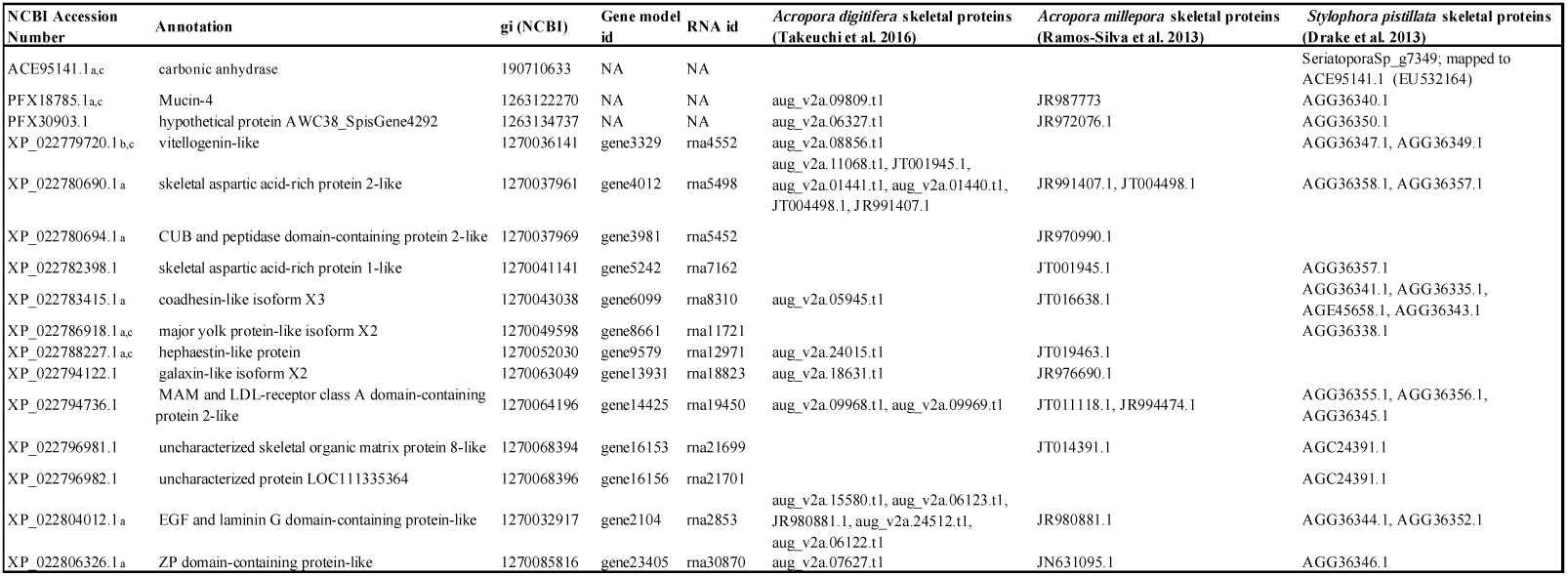
Orthologous coral skeletal proteins in the present work and previously published. Gene ontology categorization is represented as ^a^ECM/transmembrane and protein modification, ^b^membrane processing, and ^c^vesicle/secretion and metal binding.

### Skeletal proteome characterization

We interrogated the mechanisms by which proteins may be exported from or attached to the cell (SI Table 2). Seventeen coral skeletal proteins with likely complete N-terminus predictions possess signal peptides as a potential mechanism for export from the cell. Eight proteins contain at least one transmembrane span suggesting that they are embedded in the cell membrane. Further, 10 proteins likely interact with the exterior of the cell membrane by GPI anchors. In total, 25 of the 60 sequenced coral skeletal proteins exhibit documented characteristics for localization in the ECM.

Because the majority of the proteins sequenced from the *S. pistillata* skeleton do not possess features for signaling their export from the cell, we queried the data set for further suggestions of positioning the proteins in the membrane or that the proteins may be exported by vesicles such as those that may be involved in calcium concentration. To do this, we examined the skeletal proteome annotations and GO classifications toward finding common features to allow grouping of proteins. Out of the entire skeletal proteome sequenced in this study (60 proteins), 39 genes were returned with GO terms that allowed their classification into five groups of interest based on their cellular component, biological process, and molecular function to suggest likely cellular locations pertinent to the calcification mechanism, which may therefore be indicative of their function in this process: lipid\phosphate\glycan related proteins (i.e., membrane processing); ECM-related, transmembrane, and protein modification proteins; and metal binding proteins vesicular/secretion related proteins, (SI Table 3).

Of the proteins with GO terms, 10 proteins are suggested to be involved with processing of the cell membrane (Figure 3), A much larger number are related to vesicles/secretion as well as binding metal, with 21 and 25 assigned to each category, respectively. We combined these two categories in our proposed cellular location Figure 3, as some of the skeletal proteins proposed to be found intracellularly in vesicles are also known to bind calcium [34, 62]. Finally, 19 and 12 are potential ECM proteins or are involved in protein modification, respectively. Of the proteins in the vesicles/secretion and ECM categories, several appear to be completely predicted, based on comparison to orthologs in the *A. digitifera* genome, yet lack a signal peptide (SI Table 2). It should be noted that many proteins are assigned to multiple categories.

**Figure 3.**
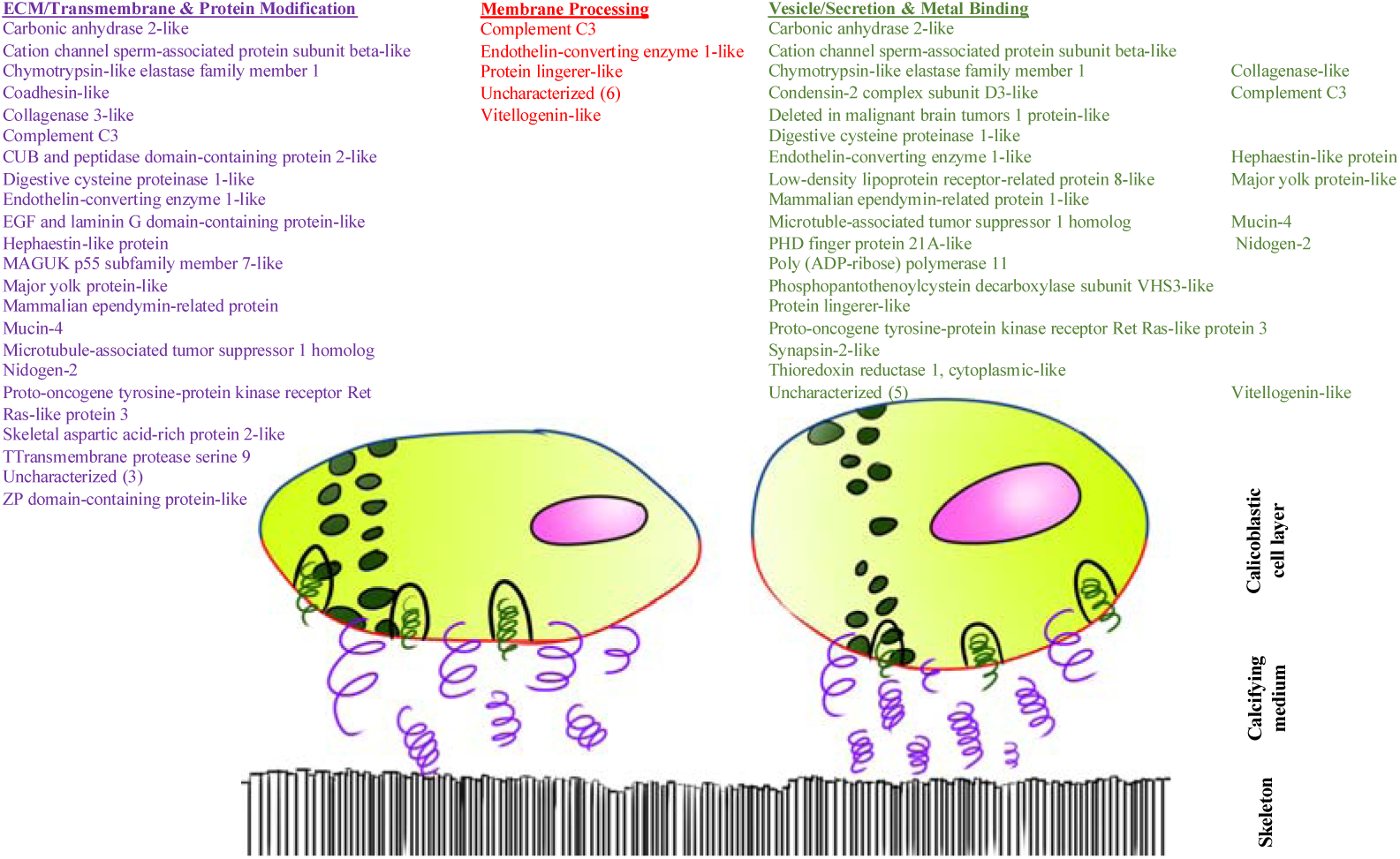
Suggested cellular and extracellular locations of *S. pistillata* skeletal protein based on GO terms. Large yellow and red circles represent calicoblastic cells with red cell membrane facing the skeleton. Small green circles represent vesicles. And purple squiggles represent transmembrane proteins that may or may not span the width of the calicoblastic space plus ECM proteins.

## Discussion

In this study we show the importance of using complementary post-extraction methods to purify and concentrate coral skeletal proteins for sequencing the full breadth of the skeletal proteome. Our results show a clear and marked difference in detected proteins between protein purification methods, with only two proteins observed in all methods. Centrifugal Filter (CF) methods yielded a much greater abundance and diversity of proteins than did acetone precipitation (ACT) (Figure 2). Moreover, we found that protein detection is largely method-exclusive (Figure 2). We hypothesize that the protein yield differences between purification methods is based on the different properties of each method. CF is a mechanical filtration based on size and has a bias toward hydrophobic proteins, whereas ACT is based mainly on the chemical interactions of proteins resulting in increased precipitation of hydrophilic proteins [63]. We speculate that the CF methods yielded more total proteins in our study as acetone precipitation does not recover all proteins [64] and those proteins which are recovered in a pellet may be difficult to resolubilize due to molecular interactions formed within the aggregate by protein denaturation [65], potentially leading to the loss of many proteins in the pellet which are not transferred to the trypsin digestion step.

A major challenge in working with intra-skeletal proteins is isolating the true skeletal proteins from soft tissue contamination [40, 41]. In the present study, as in previous work on *S. pistillata* skeletal proteins [40] we carried out an intensive oxidative cleaning step on the skeletal powder, in addition to cleaning the skeletal fragments, to avoid contamination. We did not observe any organic residues on cleaned powders examined by SEM (SI Figure 1a,b) or in PBS soaked on the powders and concentrated (SI Figure 1c), and we are therefore confident that all sequenced proteins are endogenous to the skeleton and are not coral cellular contamination.

In this study, we examined the protein composition of the SSOM and ISOM proteins separately. Although we see variations due to the different purification techniques (Figure 2), our results indicate that, broadly, the ISOM is distinguished in composition from the SSOM. Across all purification techniques, the ISOM fractions yielded more total proteins and fraction-exclusive proteins than the SSOM fraction did (Figure 2B, C). Since all extraction and purification experiments were carried out simultaneously, we can rule out batch effect and technical differences. This ISOM versus SSOM skeletome fractional composition is similar to previous studies carried out on other marine biomineralizers such as mollusks [45] as well as to the stony corals *A. millepora* [32] and *S. pistillata* [31]. However, the overlap between the two solubility fractions reported for *A. millepora* [32] was 61%, in contrast to our study which shows an overlap across all purification treatments combined of only 10% (Figure 2). One plausible explanation for this difference is the use of improved LC-MS/MS performance over the past several years enabled us to detect more proteins and obtain a more comprehensive proteome. In previous studies of coral skeletal proteomes which detailed differences between solubility fractions, the instruments used were low resolution, low mass accuracy, which tend to result in a lower percent identification of the data. Additionally, most search engines and FDR-calculating algorithms struggle with very small datasets [66, 67]. Our use of Byonic helps to alleviate statistical limitations that can result in false negative results.

Further differences in proteomes, beyond species differences, are the differing reagents used in precipitation (compared to [31]) and our smaller centrifugal filter cutoffs (compared to [32]). This second difference is particularly important with respect to protein degradation. Even when embedded in biominerals so that amino acids and even short peptides persist, proteins may still succumb to degradation [68]. This result in small peptide fragments that may be lost from centrifugal filter units with 10 kDa pore sizes and larger. It is reasonable to assume that coral skeletal proteins go through the same process, and if so, the cutoff of the membrane directly affects the number of peptide spectra matches (PSM). SDS-PAGE analysis shows smearing of extracted skeletal proteins, which in addition to likely differential addition of glycans, sulfates, and phosphates to proteins [e.x., 30, 69, 70], may indicate protein degradation (SI Figure 2). Using smaller cutoff filters in this study might have allowed us to capture some of these sheered peptides and led to higher PSMs.

### Traditional protein trafficking by signal peptides and transmembrane domain does not explain the full extent of protein transportation to skeleton

Corals’ skeleton is external to the animal; therefore, proteins in the skeletal matrices must be transported outside the cells or span the membrane and have an extracellular portion to reach the skeletal crystallization front. Indeed, recent studies of anthozoans reveal a significant proportion of TM domain proteins (∼35%) in the SOM [32, 33], similar to that found in the better-studied Echinoidia [71]. Based on these findings, we examined the hypothesis that many of the skeletal proteins originate in the plasma membrane.

Our analysis revealed that TM domain proteins are not the major component of the SOM protein complex; in the present proteome, TMHMM prediction suggests that eight are embedded in the membrane. Hence, we examined a cellular secretion option. Ramos-Silva et al. [32] reported 15 proteins (41%) with SPs that did not also possess a TM domain in the *A. millepora* proteome. In their study of the *A. digitifera* proteome, Takeuchi et al. [33] reported a similar proportion of proteins with SPs but no TM domains (40%). In the present S. *pistillata* proteome, 17 out of 60 (28%) were positive for SPs (SI Table 2, [31]), likely a combination of incomplete gene prediction and other mechanisms for exporting the proteins to the calicoblastic space.

Hydrophobic regions of TM domains are similar to those of SPs, making SP prediction difficult [54, 72]. We therefore cannot say for sure, based on these two analyses (TMHMM and SignalP) alone, which is the more dominant pathway for protein transport for many of the SOMPs. Further, while SP by itself can help predict traditional secretory pathways [72], proteins that do not possess classical SPs or TM regions may still leave the cell via other means (reviewed in [72-74]). This includes membrane pores [72], ATP-binding cassette transporters [75] and autophagosome/endosome-based secretion [73, 76], as well as exosomes [77] which can transport proteins that lack signal peptides in extracellular vesicles [78]. Since TM and SP analysis did not fully explain the mechanism of protein transportation, we classified the likely functions and locations of the skeletal proteins according to their GO terms as lipid\phosphate\glycan related proteins, metal binding proteins, vesicular/secretion related proteins, ECM-related and transmembrane proteins, and protein modification proteins (Figure 3, SI Table 3). Looking at the proposed locations or functions of skeletal proteins (Figure 3), we find comparable numbers of vesicular and metal binding proteins versus TM domain/ECM proteins. Further, several of these proteins appear to be completely predicted at their N-termini yet lack signal peptides (SI Table 2). These results are in-line with the trend reported in previous coral skeletal proteomes [31-33]. These proteins may leave the calicoblastic cells through one of the non-classical mechanisms described above. While we do not exclude the role of SP and TM proteins in the skeletal deposition process, we suggest that the biological mechanism of SOM transportation is enhanced by exosomes and other non-traditional secretion pathways.

A further method by which proteins may be exported to the site of calcification is a vesicular pathway that differs from the conventional SP and TM pathway and remains to be fully characterized. Previous studies have shown Ca^2+^ rich granules in the calicoblastic epithelium (skeletogenic cells), but not in the other tissue layers, suggesting their role as a Ca^2+^ reservoirs in the cells. Vesicles were previously identified in corals [79, 80]; however, their origin and content was not detailed, and they are sometimes attributed to preservation byproducts. These intracellular ion-rich vesicles may endocytose sea water through macropinocytosis [81, 82], after which they are enriched in carbonate ions and then form hydrated ACC and anhydrous ACC precursors stabilized by acidic biomolecules including CARPs [83]. Using cell cultures, Mass et al. [62] suggest that the vesicles, which contain Asp rich proteins, then transport their contents to the ECM, releasing their content by exocytosis. The biomineral then further develops extracellularly, likely aided by other ECM proteins [30, 35, 50, 84] as well as other biomolecules [85-88]. At present the processes of calcium delivery to the skeleton and the roles of most of the proteins in coral skeletal deposition remain to be determined.

It is difficult to map and characterize proteins in non-model organisms such *S. pistillata* [89-91]. High quality proteomic mapping requires knowledge of phosphorylation, glycosylation, proteolytic cite activities and other modifications [31, 92-94], in order to create a more thorough database [89-91]. Further, many of the coral skeletal proteins reported to date remain uncharacterized. Uncharacterized proteins were reported in *Acropora* skeletons at a rate of approximately 25% [32, 33] while, they were previously reported at less than 10% in *S. pistillata* skeleton [31].

Our study revealed a greater proportion, of 28% uncharacterized proteins, in *S. pistillata* skeleton, in line with *Acropora* spp. While partially attributed to sample size, it is most likely due to quality of genomic data available, since stony corals are non-model organisms and their genomic libraries are far from complete, resulting in incomplete databases on which to map the proteome and many uncharacterized genes.

## Conclusion

In this study we have considered the differential effects of coral skeletal protein extract preparation as well as the method by which these proteins, or parts thereof, are transported from intracellular to extracellular locations. When preparing coral skeletal proteomes, we propose that a multi-method approach to cleaning, demineralization, and protein purification should be used. Our results showed that each protein preparation protocol yielded exclusive sets of proteins with little overlap between ACT and CF fractions. While CF protocols yielded many more proteins than did ACT methods, use of a single protocol to clean and concentrate coral skeletal proteins results in a significant amount of data loss, and it is therefore of crucial importance to consider alternative and complementary methods to obtain a fully comprehensive skeletal proteome. We showed that while the role of TM domain proteins cannot be overlooked, many of the proteins detected in the *S. pistillata* skeletal proteomes as well as in that of other species point toward other secretory or vesicular pathways.

Our categorization method, supported by data from other recent studies, also suggests that corals use an alternative secretory pathway, such as exosomes or non-classical secretion vesicles, and much work is required in order to determine the calcium deposition pathway and the proteins involved. Our study provides a large set of new uncharacterized coral skeletal proteins as well as others of purported function but that have not been observed before in the coral SOM. These data expand the current knowledge of the SOM in corals and will help, in future studies, to resolve corals’ calcium deposition mechanism and the various roles of the proteins involved.

## Abbreviations

(ACT): acetone precipitation of proteins
(CF): centrifugation ultrafiltration of proteins
(FDR): false discovery rate
(GO): gene ontology
(GPI): glycosylphosphatidylinositol
(ISOM): insoluble skeletal organic matrix
(PSM): peptide spectra matches
(PBS): phosphate buffered saline
(SOM): skeletal organic matrix
(SOMP): skeletal organic matrix proteins
(SSOM): soluble skeletal organic matrix
(TM): transmembrane

## Declarations

### Ethics approval and consent to participate

Coral nubbins were collected under a special permit from the Israeli Natural Parks Authority.

### Consent for publication

Not applicable.

### Availability of data and materials

The datasets generated during the current study are available in the ProteomeXchange repository under file number PXD017891 (https://www.ebi.ac.uk/pride/archive). The reviewer login username and password have been provided to the reviewers. Data will be made publicly available without a required login in ProteomeXchange upon publication of the manuscript.

### Author contributions

YP, JD, and TM designed the study. YP and RA prepared the samples. YP, RA, AM, ML, DM, and JD analyzed the data. All authors wrote the manuscript and approve this submission.

## Acknowledgements

We thank staff at the Bioinformatics Core Unit, University of Haifa, and The Crown Genomics Institute of the Nancy and Stephen Grand Israel National Center for Personalized Medicine, Weizmann Institute of Science.

