## Supplemental Information for "Optimization of skeletal protein preparation for LC-MS/MS sequencing yields additional coral skeletal proteins in *Stylophora pistillata*"

|  | Coral Species | Reference | Cleaning steps | Demineralization | Extraction procedure |
| --- | --- | --- | --- | --- | --- |
|  | *Acropora millepora* | Ramos-Silva et al., 2013b, 2013a | (1) Washed fragments in  NaOCl 5% v/v, 72 h.  (2) Powder (*<*200 μm) in  NaOCl 1% v/v, 5 h. | Acetic acid 10% v/v overnight at 4^0^C  until pH 4 | ASM: Centrifugation Ultrafiltration Dialysis |
|  |  | Ramos-Silva et al., 2013a |  |  | AIM: 6 × Centrifugation with Milli-Q water |
|  | *Acropora digitifera* | Takeuchi et al., 2016 | (1) Washed fragments in NaClO 12.5%v/v and water  (2) powder fragments in NaClO 12.5%v/v and water | Acetic acid 1M 24h | ASM: Chloroform/ methanol precipitation |
|  |  |  |  |  | AIM: Water wash, solubilization buffer (1% SDS, 10mMDTT, 50mMTris-HCl (pH 8.0)) |
|  | *Stylophora pistillata* | This study | (1) Washed fragments in 1:1 NaOCl 3%:H_2_O_2_ 30% under sonication 1h incubation 24h at R.T + milliQ wash X5 (2) Powder <63μm, same as in (1), repeated 3 times. | Acetic acid 0.5M 4h at R.T until complete dissolution. | ASM & AIM:  Lyophilization, Centrifugation  Ultrafiltration Dialysis. 100% followed by 80% acetone precipitation |
|  | *Stylophora pistillata* | Drake et al., 2013 | (1) Washed fragments in  NaOCl 3% wt/v, 4 h.  (2) Powder (*<*150 _m)  Second bleaching. | HCl 1N at R.T pH 7 | ASM: Centrifugation |
|  |  |  |  |  | AIM: Acetone 90%  Centrifugation |

SI Table 1. Adapted from [Marie, 2013] and edited. A brief summary of different cnidaria; cleaning, demineralization and extraction protocols for skeletal matrix proteins. (ASM, Acid soluble matrix; AIM, Acid insoluble matrix; R.T, room temperature)

**File Name ‘SI Table 2.xlsx’**

SI Table 3. Byonic output of *S. pistillata* skeletal proteins sequenced by LC-MS/MS across all solubility groups and purification methods.

**File Name ‘SI Table 3.xlsx’**

SI Table 2. Proteins sequenced from *S. pistillata* skeleton grouped by gene ontology cellular location and function. A given protein may be assigned to more than one group.

A	B

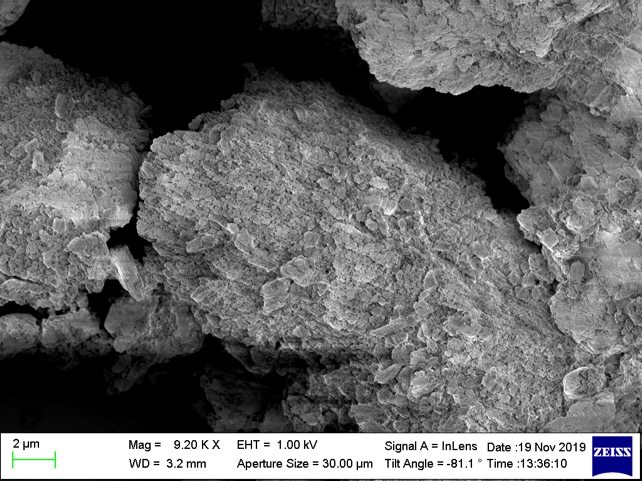

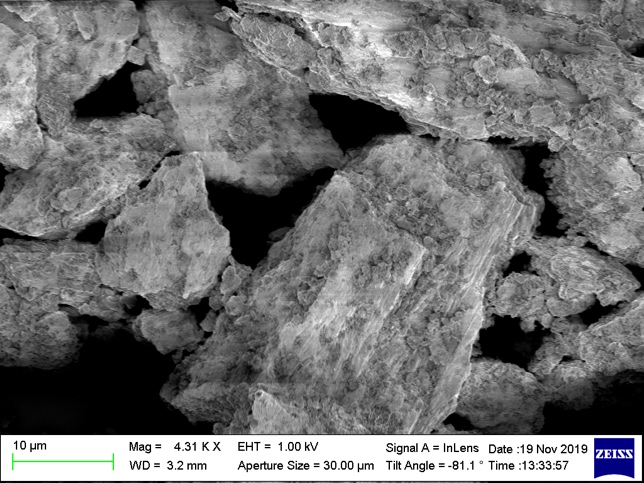

C

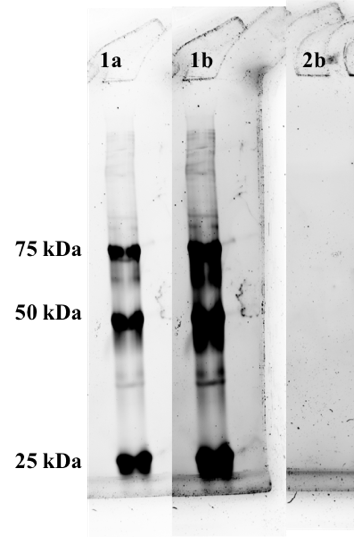

SI Figure 1. Scanning electron micrographs (a, b) showing that *S. pistillata* ground to 63 μm and cleaned as described are free of visible exogenous contaminating organic material. Soaking the cleaned ground skeleton in PBS while sonicating for an hour also did not reveal any protein banding on TGX Stain-free gels (c). Lanes in (c) are: 1a = Precision Plus Unstained ladder at auto-exposure (~5 seconds); 1b = the same lane as (1a) but exposed for 20 seconds; 1c = concentrated PBS soak exposed at 20 seconds; gels were activated under UV light for five minutes. No protein smearing or banding was observed in the concentrated PBS soak of the cleaned skeleton powder.

A	B

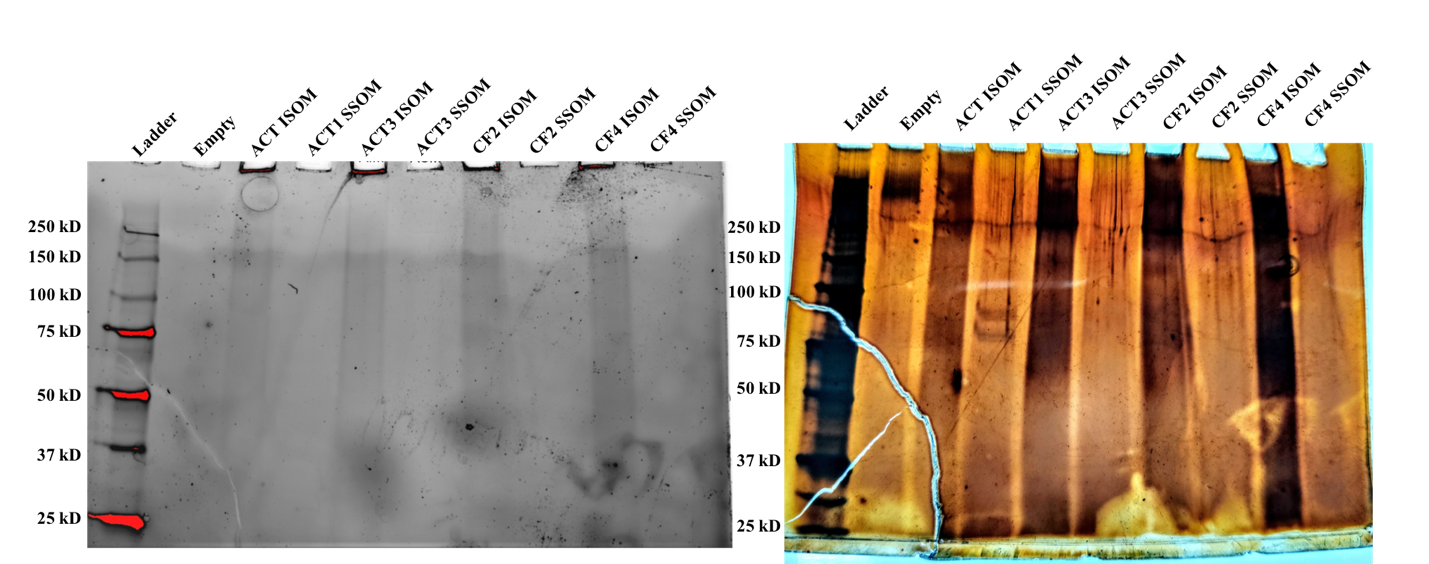

SI Figure 2. Protein gel of organic matrix proteins extracted from cleaned *S. pistillata* skeleton powders following UV activation for five minutes (A) and then silver staining (B).
